# Spatial transcriptomics identifies the *Pseudomonas aeruginosa* major outer membrane protein OprF as a critical factor in blinding corneal infections

**DOI:** 10.64898/2026.05.20.726515

**Authors:** Serena Abbondante, Hao Zhou, Nicole Chumbler, Oscar Negron, Michaela Marshall, Ankush Tyagi, Arne Rietsch, Eric Pearlman, Mihaela Gadjeva

## Abstract

*Pseudomonas aeruginosa* is a globally recognized pathogen causing pulmonary, skin, and severe corneal infections (keratitis), with the potential to induce irreversible blindness if untreated. Spatial transcriptomic analysis of *P. aeruginosa* infected corneas identified elevated expression of the outer membrane proteins OprF and OprL and PA1414, which encodes the small RNA SicX in the corneal stroma compared with corneal epithelium. Comparative spatial transcriptomics analysis of corneas infected with an *oprF* transposon (TN) mutant showed reduced expression of the type III effector protein ExoT, which was absent in an *oprF* deficient mutant (Δ*oprF*) and in contrast to PA14, did not inhibit reactive oxygen species (ROS) production by neutrophils. Corneal infection with the Δ*oprF* mutant resulted in reduced corneal virulence and lower CFU compared to the parental PA14 strain. Collectively, our findings demonstrate a coordinated virulence program connecting OprF functionality with the release of ExoT and its ability to block ROS production and survive in infected corneas.

## INTRODUCTION

*Pseudomonas aeruginosa* is a major cause of pulmonary and wound infections, and is also an important cause of blinding corneal infections that are prevalent in the USA and worldwide (1). Patients with corneal ulcers caused by *P. aeruginosa* are highly enriched in neutrophils that control bacterial replication by oxidative and non-oxidative mechanisms, and release serine and matrix metalloproteinases that digest the stroma, resulting in corneal opacification and visual impairment. While antibiotics remain the first line of treatment, there are increasing reports of antibiotic resistance among clinical isolates of *P. aeruginosa*, including a 2023 outbreak of drug-resistant *P. aeruginosa* keratitis linked to contaminated artificial tears that resulted in blinding disease (2, 3). Approximately 30% of *P. aeruginosa* clinical isolates exhibit multidrug resistance (4); therefore, the increased difficulty in treating *P. aeruginosa* infections highlights an unmet need to develop new anti-bacterial therapies, potentially by targeting critical bacterial proteins.

*P. aeruginosa* expresses multiple virulence factors, including type III secretion exoenzymes and Type IV pili that have an important role in corneal infections (5–8). Porins, which are outer membrane proteins involved in functions that include solute transport and iron uptake, are also required for virulence in a murine lung infection model (8). We recently described an unbiased spatial transcriptomics approach to characterize bacterial gene expression in *P. aeruginosa* infected murine corneas (9). That study identified several bacterial genes that were upregulated, including a previously uncharacterized gene, PA2590, which shows structural homology with *E.coli* pyochelin iron transporter that can potentially bind the pyochelin siderophore (9). Furthermore, corneal infection with a PA2590 transposon mutant resulted in significantly impaired bacterial growth and less severe disease (9), thereby identifying PA2590 as a putative *P. aeruginosa* virulence factor.

In the current study, we performed an independent spatial transcriptomics analysis and showed elevated expression of several genes in the corneal stroma compared with the corneal epithelium, including the major outer membrane proteins OprF. We generated Δ*oprF* deletion mutants and demonstrated that OprF is required for expression of the type III effector ExoT, which regulates production of reactive oxygen species (ROS) by neutrophils and that OprF is required for full virulence in infected corneas.

## RESULTS

### Spatial transcriptomics reveals differential enrichment of OprF, OprL, and PA1414 in *P. aeruginosa* infected corneas

To examine expression of potential *P. aeruginosa* virulence factors in a clinically relevant model of infection, we used a well-established murine model of blinding keratitis where corneas were infected topically with 5 x 10^5^ log phase *P. aeruginosa* strain PA14 and outcome is measured by disease severity and recovery of viable bacteria (5, 10).

Twenty-four hours after infection, eyes were sectioned by cryostat and processed for spatial transcriptomics analysis using *P. aeruginosa* specific probes (Nanostring, **Figure S1B**). This panel is distinct from the one used in our prior study (9) and includes probes targeting the major outer membrane protein OprF, the outer membrane lipoprotein OprL, the recently identified virulence factor PA1414 (11) and Type III secretion system genes, including ExoT.

Spatial expression of the probed 20 bacterial transcripts in the infected tissue showed that all transcripts were detected in at least one region of interest. Combined analysis of regions of interest revealed increased transcript abundance in the corneal epithelium and stroma relative to the anterior chamber, including housekeeping ribosomal genes *rpsA* and *rplL* and the ribose-5-phosphate isomerase gene *rpiA* (**Figure 1A)**. Notably, the normalized transcript abundance of *oprL*, *oprF*, and PA1414 was enriched in the infected stroma compared with the epithelium (**Figure 1B, C)**.

**Figure 1.**
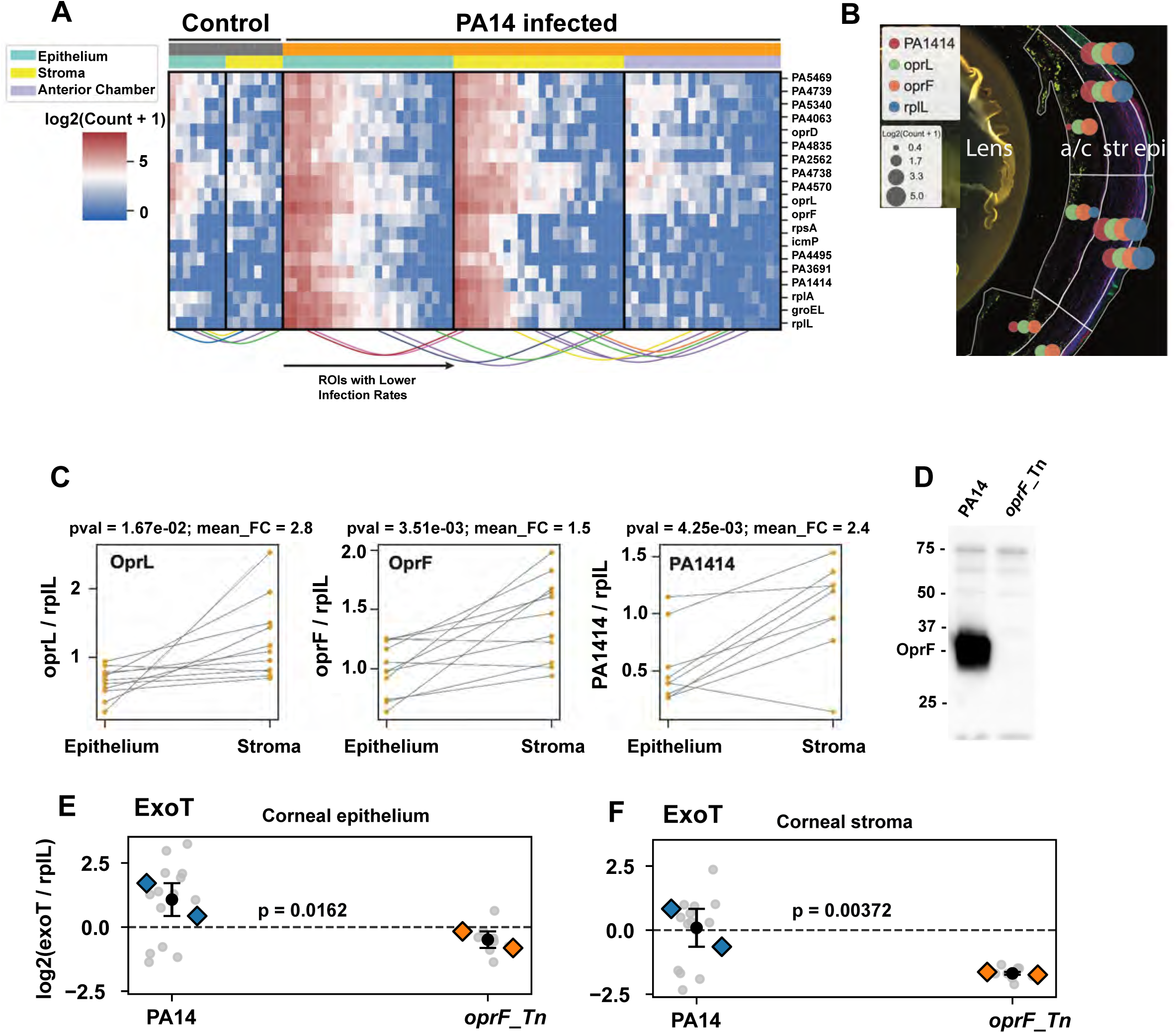
Spatial transcriptomics of PA14 infected corneas. Formalin fixed sections of PA14 infected corneas were incubated with a panel of 20 probes for *P. aeruginosa* genes using the GeoMx Digital Profiler platform. **A.** Gene expression in the corneal epithelium (green), stroma (yellow) and anterior chamber (purple). Each column represents an individual Region of Interest (ROI). Levels for 20 transcripts are shown for 22 ROIs at each site. Colored lines at the bottom connect neighboring ROIs. Data are presented cumulatively across technical of 2 biological replicates. **B.** Representative image of *P. aeruginosa*-infected eye cross-section stained for cytokeratin (green), Ly6G neutrophils (magenta), and DNA (blue) visualized using GeoMx DSP. Logarithmic transcript abundances (log2(count + 1)) shown in the table (circle size) for OprL, OprF, PA1414, and rplL housekeeping genes in infected corneal epithelium, stroma, and anterior chamber are represented as circles. **C.** Relative expression of *oprL*, *oprF and PA1414* transcripts in the corneal epithelium and stroma are normalized to the housekeeping *rplL*. Data points represent individual ROIs; FDR values and fold change (FC) are noted for each gene. **D.** Absence of OprF protein in the PA14 *oprF::Tn* mutant. **E, F.** ExoT expression in corneal epithelium € and stroma (**F**) of C57BL/6 mice infected with PA14 WT or the PA14 *oprF::Tn* mutant. Data are shown as log₂(exoT/rplL) (rplL housekeeping gene) based on GeoMx regions of interest (ROIs) (gray data points), N=2 mice (blue and orange data points). P values were derived from a linear mixed-effects model using ROI values per mouse as a random intercept (log2(exoT/rplL).

Furthermore, spatial transcriptomic analysis of corneas infected with PA14 or the OprF transposon mutant *orpF::Tn* revealed that ExoT expression was significantly reduced in both the stroma and epithelium of *orpF::Tn* infected corneas **(Figure 1D-E)**. In contrast, expression of the translocon proteins *popB* and *popD* remained unchanged (**Figure S1A)**. These and other differences in gene expression between PA14 and *orpF::Tn* were also found using the larger transcript library from our previous study (**Figure S1B**) (12). Collectively, these findings indicate transcriptional reprogramming of *P. aeruginosa* in distinct corneal microenvironments during infection, and also indicate a requirement for OprF in expression of ExoT.

### OprF mutants induce increased ROS production by neutrophils

To investigate the role of OprF in bacterial pathogenesis, we generated *oprF* deletion mutants (Δ*oprF*) using primers targeting the promoter region of the gene as described in the Methods. Complemented strains were generated by reintroducing the *oprF* gene into the mutant background on a plasmid. Independent clones of both the deletion mutant and the complemented strain exhibited growth rates comparable to the parent strain *in vitro* (**Supplementary Figure 2A**). Consistent with our spatial transcriptomics analysis showing reduced *exoT* transcript abundance in the Δ*oprF* mutant, ExoT protein was detected in PA14 and the complemented Δ*oprF*:*oprF* strain, but was absent in the Δ*oprF* mutant (**Figure 2A,B**), indicating that OprF is required for ExoT expression.

**Figure 2.**
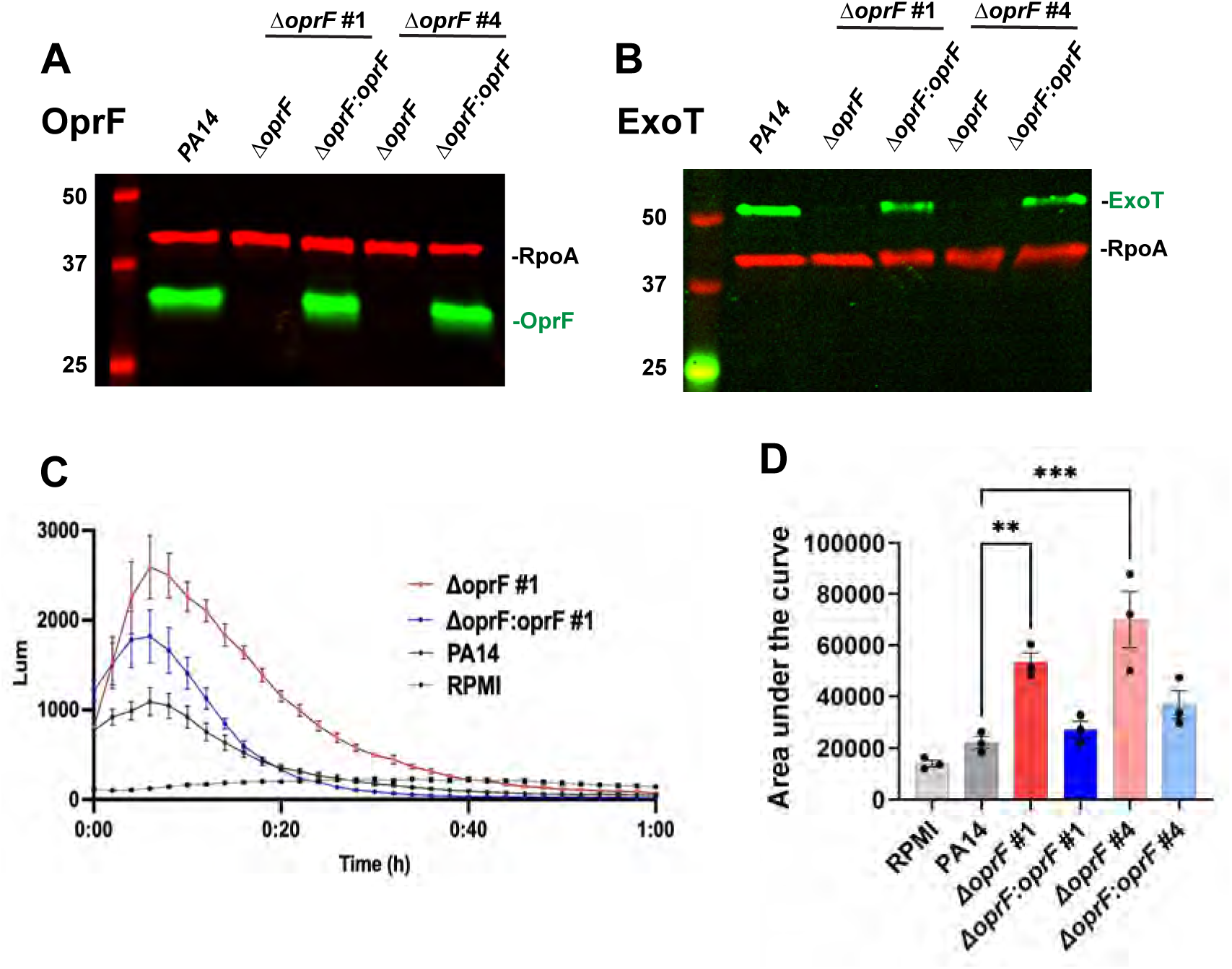
The role of OprF in Reactive oxygen species (ROS) production. **A.** OprF protein and housekeeping RpoA in PA14, Δ*oprF or* Δ*oprF:oprF* (complemented) clones #1, and #4. **B.** ExoT protein in Δ*oprF* and Δ*oprF:oprF* clones. **C**. Time course of ROS production by bone marrow neutrophils infected with PA14, Δ*oprF or* Δ*oprF:oprF* detected by Luminol (mean +/- SEM). **D.** Total ROS production over 1h (mean +/- SEM of area under the curve of the time course calculated using GraphPad Prism. N=3 biological replicates. Statistical significance was determined using 2-way ANOVA analysis with Tukey’s multiple comparisons. **: p<0.01; ***: p<0.001. Experiments were repeated x3.

We reported that neutrophil production of reactive oxygen species (ROS) is essential to kill *P. aeruginosa,* and that bacteria expressing type III secretion system (T3SS) exoenzymes ExoS or ExoT survive by blocking assembly of NADPH oxidase proteins and inhibiting ROS production (13). We therefore examined ROS production by neutrophils infected with PA14, the Δ*oprF* mutant and the complemented strain (Δ*oprF*:*oprF*). ROS production was quantified using the Luminol assay.

We found that PA14-infected neutrophils induced only a transient and low level of ROS, consistent with our earlier findings using strain PAO1 (13). In contrast, neutrophils infected with the Δ*oprF* mutants #1 and #4 exhibited significantly increased production of reactive oxygen species (**Figure 2C-D**, time course of #4 shown in **Figure S2B**). As expected, ROS production by the complemented Δ*oprF:oprF* strain was comparable to that induced by the parental strain. We also found that ROS production was inhibited by the NADPH oxidase inhibitor diphenyleneiodonium chloride (DPI) (**Figure S2C**).

Collectively, these findings demonstrate that OprF is required to suppress ROS production in PA14 infected neutrophils, and that increased ROS production in ΔOprF infected neutrophils is associated with loss of ExoT.

### OprF is required for *P. aeruginosa* virulence in infected corneas

OprF is the major outer membrane protein and the most abundant non-lipoprotein component of the outer membrane, contributing to the structural integrity of the bacterial cell envelope in addition to biofilm formation, motility, and virulence (14–17). Given the observed spatial enrichment of *oprF* transcripts in the corneal stroma and the role of OprF in ROS production, we next examined if there is a role for OprF in *P. aeruginosa* virulence following corneal infection. The corneal epithelium of C57BL/6 mice was abraded with 3 parallel 3mm scratches, and infected topically with 5 x 10^5^ log phase *P. aeruginosa* strain PA14 or the PA14 *oprF* deletion mutant (Δ*oprF*). After 48h, mice were euthanized, corneas were imaged, and viable bacteria were quantified by colony-forming units (CFU) following homogenization of whole eyes.

Healthy corneas are transparent, in part due to the absence of inflammatory cells. However, following infection with PA14, there was pronounced corneal opacification (**Figure 3A**). In contrast, mice infected with the Δ*oprF* mutant exhibited significantly reduced corneal opacity compared to the parental strain (**Figure 3A, B**). Consistent with this finding, Δ*oprF*-infected corneas contained significantly fewer viable bacteria, as measured by colony-forming units (CFU), compared to PA14 infected corneas (**Figure 3C**).

**Figure 3.**
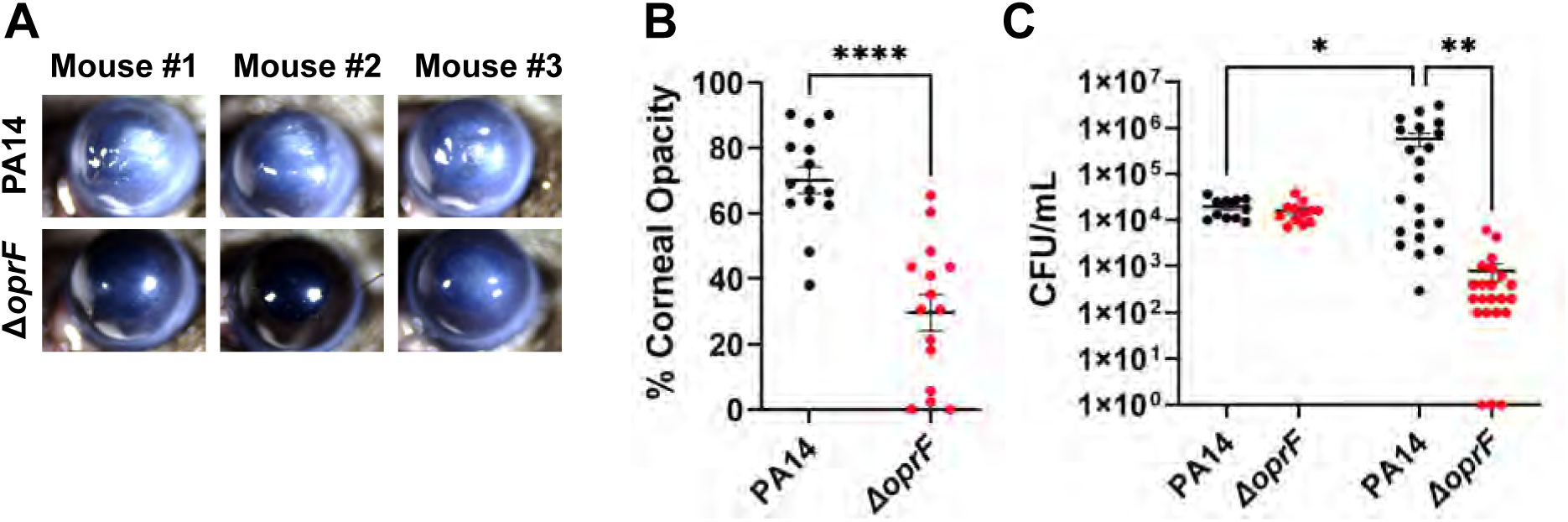
Role of OprF in *P. aeruginosa* corneal infection. Corneas of C57BL/6 mice infected with 5×10^5^ CFU PA14 or PA14 *ΔoprF* clone 1,. **A.** Representative images of infected corneas after 48h; **B.** Corneal opacification quantified using Image J and **C)** viable bacteria by CFU. Each data point represents a single infected cornea and data show three experiments combined. Statistical significance was determined using 2-way ANOVA analysis with Tukey’s multiple comparisons. *: p<0.05; **: p<0.01; ***: p<0.001; ****: p<0.0001.

Taken together, these findings identify a critical role for OprF in the ability of *P. aeruginosa* strain PA14 to replicate in the cornea and cause disease.

### Impaired neutrophil recruitment to corneas infected with Δ*oprF* mutants

In this model of *P. aeruginosa,* we reported that neutrophils comprise >80% of the total CD45+ cellular infiltrate to the corneal stroma in the first 48-72h, with inflammatory monocytes comprising most of the remaining CD45+ infiltrating cells (5, 10). To characterize the cellular response in corneas infected with the Δ*oprF* mutant, mice were infected with *P. aeruginosa* strain PA14 or Δ*oprF*, eyes were fixed and paraffin-embedded, and corneal sections were immunostained with antibodies to OprF and with antibodies to Ly6G that identify neutrophils. Sections were also stained with DAPI to detect total cell nuclei.

OprF-expressing bacteria were readily detected in the PA14, but not in Δ*oprF* infected corneas, and co-localized with Ly6G+ neutrophils (**Figure 4A**). To quantify neutrophil and monocyte infiltration, infected corneas were digested with collagenase and total CD45+ myeloid cells were isolated and incubated with antibodies for quantification of live cells by flow cytometry analysis. Neutrophils were defined as CD45⁺/Ly6G⁺/CD11b⁺ and monocytes as CD45⁺/Ly6G⁻/Ly6C⁺/CD11b⁺. A representative scatter plot shows neutrophils comprising ∼93% total infiltrating cells, whereas monocytes were ∼3.5% in PA14-infected corneas (**Figure 4B**) Gating strategy is shown in Supplementary Figure 4.

**Figure 4.**
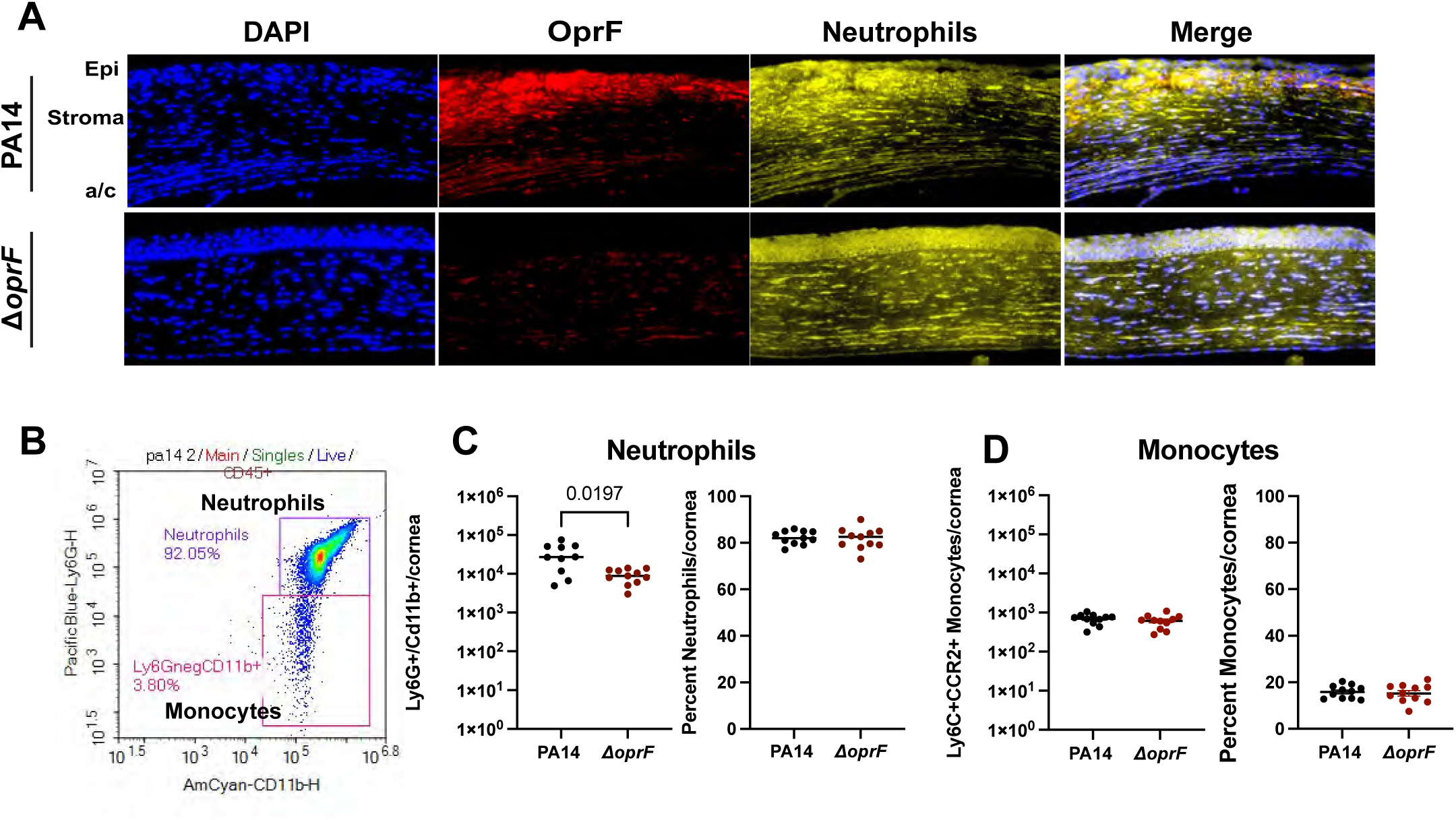
Role of OprF in neutrophil infiltration to infected corneas. Corneal sections showing OprF and Ly6G+ neutrophils in the corneal stroma and epithelium 24h after PA14 infection. DAPI staining indicates cell nuclei. **B.** Representative scatter plot of live, single, CD45+ cells identifying Ly6G +, CD11b+ neutrophils and Ly6C +, CD11b+ monocytes 24h post infection. **C.** Total and percent neutrophils and monocytes 24 h after corneal infection with PA14 or *ΔoprF*. Gating strategy is shown in Supplemental Figure 4. Data points represent individual infected corneas and data are representative of three independent experiments. Statistical significance is based on 2-way ANOVA analysis with Tukey’s multiple comparisons. *: p<0.05; **: p<0.01.

The total number of neutrophils in infected corneas was significantly lower in Δ*oprF* compared with PA14 infected corneas although there was no difference in percent (**Figure 4C**). There was no difference in total or percent monocytes in corneas infected with PA14 or Δ*oprF* (**Figure 4D**). These findings indicate that the reduced corneal disease severity observed in mice infected with the Δ*oprF* mutant reflects decreased neutrophil recruitment to infected corneas, which is also consistent with fewer bacteria.

Collectively, these findings demonstrate that OprF is required to protect *P. aeruginosa* strain PA14 from neutrophil killing in infected corneas and to cause the corneal opacification associated with visual impairment.

## DISCUSSION

Spatial transcriptome analysis preserves tissue architecture and provides spatial context, enabling identification of distinct gene expression patterns and revealing unique insights into the spatial dynamics of infection. Using this approach, we identified three *P. aeruginosa* transcripts that were elevated in the stroma compared with the epithelium: PA1414, which encodes *sicX* and is induced under anaerobic conditions (11), consistent with the relatively anaerobic environment of the inflamed corneal stroma (18). We also found elevated expression of the lipoprotein precursor gene *oprL* (19), and *oprF*, which encodes the major outer membrane protein OprF (20).

OprF is the most abundant non-lipoprotein component of *P. aeruginosa* outer membrane and functions to maintain the structural integrity of the bacterial cell envelope. It is also implicated in biofilm formation, motility, and virulence (14–17). Structurally, OprF comprises eight antiparallel beta sheets forming a beta-barrel and a flexible periplasmic domain. OprF plays critical roles in maintaining cell shape, regulating osmotic balance, and facilitating the passage of large solutes up to 3 kDa in size. Furthermore, OprF interacts with the bacterial biofilm matrix through a LecB-Psl interaction, forming a protein-mediated lipid-exopolysaccharide complex that facilitates stability of the biofilm matrix (21). OprF is also a major component of bacterial outer membrane vesicles (OMV), which transport signaling molecules that mediate bacterial communication with other bacteria and with host cells (22). Loss of OprF promotes increased OMV generation attributed to the 4-fold increase in production of the quorum-sensing *Pseudomonas* quinolone signals (PQS) (23). The impaired adhesion of the OprF-deficient strains to epithelial cells, dysregulated OMV shedding, and decreased biofilm formation may contribute to the attenuated virulence observed in the strains lacking OprF in several infection models (24, 25). OprF is also reported to bind IFNγ, which induces expression of a quorum sensing protein in the bacteria (26).

We examined the role of OprF in the virulence of *P. aeruginosa* in blinding corneal infections and found that deletion of *oprF* resulted in rapid clearance from infected corneas and less severe corneal disease than infection with the parent PA14 strain. *In vitro*, we demonstrated that *oprF* mutants induced ROS production by infected neutrophils, indicating that OprF is required to suppress NADPH oxidase activity in these cells. As we previously reported that type III secretion exoenzymes ExoS and ExoT actively inhibit NADPH oxidase assembly, we examined ExoT production in Δ*oprF* mutants (PA14 does not produce ExoS). Our finding that ExoT protein is absent in Δ*oprF* mutants likely explains the impaired ability of these mutants to inhibit ROS production by neutrophils *in vitro* and is consistent with increased susceptibility to neutrophil killing in infected corneas. These findings also indicate a potential regulatory role for OprF in expression of ExoT, although it is not clear if this is a direct effect on ExoT or more likely an indirect role of OprF given the multiple effects of this outer membrane protein on *P. aeruginosa* physiology (16, 27).

The ability of microbes to survive *in vivo* depends on their resistance to being killed by host immune cells, especially neutrophils that have evolved robust antimicrobial mechanisms. *P. aeruginosa* expresses multiple virulence factors that promote bacterial survival after phagocytosis by neutrophils. While neutrophils utilize non-oxidative mechanisms to control microbial growth, including anti-microbial peptides and nutritional immunity through iron and zinc sequestration that starve microbes of basic elements (28), NADPH oxidase plays a major role in killing ingested microbes. To evade this microbial killing mechanism, *P. aeruginosa* expresses multiple virulence factors, including the T3SS effectors ExoS and ExoT. We previously reported that ExoS and ExoT in strain PAO1 inhibit ROS production by human neutrophils, and that ExoS ADP ribosyltransferase activity targets the Ras small GTPase, thereby blocking the assembly of the NADPH complex (13).

In contrast to the ExoT-expressing parent strain PA14, we show here that the *oprF* transposon and Δ*oprF* deletion mutants induce robust ROS production by neutrophils. Further, ExoT was absent in the Δ*oprF* mutant, thereby indicating that ExoT expression is regulated, at least in part, by OprF.

ExoT has GTPase-activating protein (GAP) activity, which contributes to immune evasion by disrupting cytoskeletal dynamics and promoting host cell death during corneal infections (29, 30). In epithelial cells, ExoT inhibits wound healing and cellular repair mechanisms by blocking cellular proliferation and spreading (31). Moreover, *P. aeruginosa can* establish an intracellular niche in corneal epithelial cells, where ExoT supports bacterial persistence. Collectively, these data highlight the importance of ExoT as key mediator of *P. aeruginosa* virulence (32). This finding is consistent with a previous report documenting reduced ExoT release in the OprF-deficient *P. aeruginosa* H636 strain (33). Consistently, the spatial transcriptomics analysis of tissues infected with the *oprF::Tn* mutant showed that the stromal corneal segments had lower ExoT transcript abundance.

In contrast to ExoT, we did not observe differences in the transcripts encoding T3SS translocator proteins *PopB* and *PopD*. While we know that expression of *ExoT* and other exoenzymes is tightly regulated and distinct from the regulation of *PopB* and *PopD*, further studies are required to identify the mechanisms by which OprF regulates ExoT expression. It is possible that the envelope stress caused by the loss of OprF promotes expression of the small RNAs such as *rsmZ*, thereby interfering with *RsmA*-driven responses such as production of type III secretion effectors (34). Deletion of *oprF* has also results in an increase in c-di-GMP (34), which in turn leads to an increase in *rsmZ* expression through an unknown mechanism (35).

Our studies therefore not only identify OprF as having an important role in a clinically relevant model of blinding corneal infection, but also indicate that this is at least partly dependent on ExoT expression. The underlying mechanisms of OprF regulation of ExoT and other genes will be the subject of future studies.

Our findings have clear translational relevance. OprF has been explored for more than a decade as a vaccine antigen and biologic target. Early clinical studies of the recombinant OprF/I (IC43) vaccine showed strong immunogenicity and good safety profiles but did not reduce infection rates in vaccinated subjects (36). Preclinical work has similarly demonstrated that immunization with OprF or OprF/OprI elicits robust opsonophagocytic antibodies in animal models, yet these approaches have not translated into effective sterilizing immunity (37). These studies examined whether the antibodies elicited opsonophagocytic responses rather than functional impairment of other virulence pathways. Our data suggest that drugs targeting OprF may function not only through opsonization, but also by disrupting OprF function, inducing envelope stress, and indirectly suppressing T3SS activity. Thus, our work identifies additional possible mechanisms of action for OprF-targeting small molecules, biologics, or vaccines, providing a new framework for drug development to overcome limitations of previously tested OprF-based vaccine candidates, which failed to achieve sterilizing immunity and showed limited clinical efficacy.

In conclusion, findings from the current study identify the major outer membrane protein OprF as having a critical role in virulence in blinding *P. aeruginosa* keratitis and show that a major function of OprF is regulation of the type III secretion system protein ExoT and inhibition of neutrophil ROS production as at least one mechanism by which OprF regulates *P. aeruginosa* virulence.

## METHODS

### Bacterial strains

*Pseudomonas aeruginosa* PA14 mutants and their sources are shown in **Table 1**. The transposon mutants were generated as a part of a transposon insertion library described by Liberati and co-workers (38).

**Table 1.**
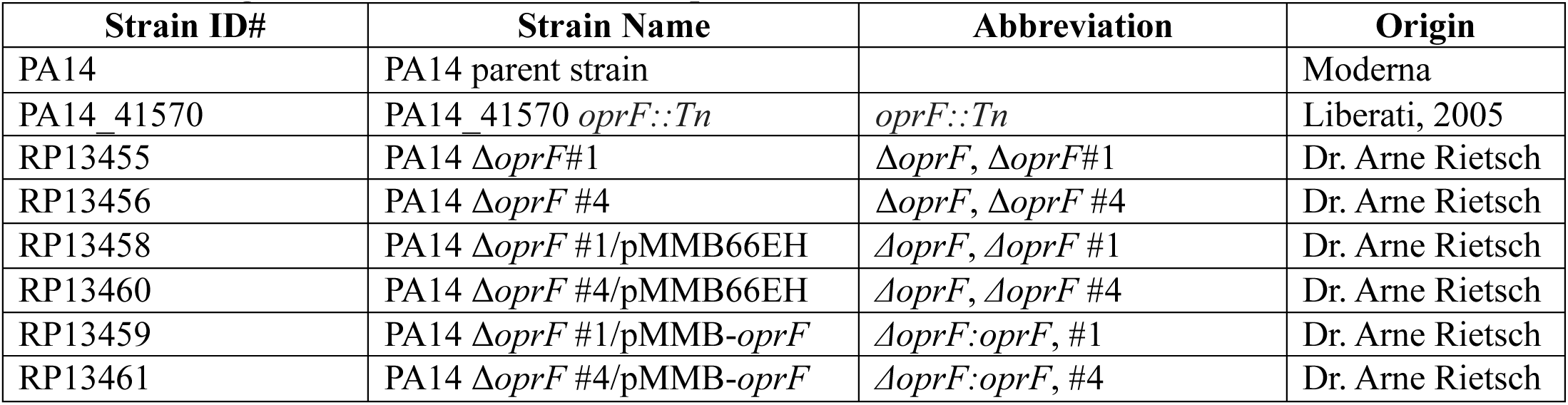
*P. aeruginosa* PA14 mutant and complemented strains.

### OprF deletion mutants and complemented strains

#### Deletion mutant

The start site of *oprF,* including potential alternative start codons (figure), was deleted by amplifying flanking DNA that specifies the extent of the deletion using PA14 chromosomal DNA and primers: oprF-5-1G ACGATCATGCGCACCCGTGGAAATTAATTAAGGTACCGGTTGAAGAACTTTGAGGGC AAG-3’) and oprF-5-2 (5’-GTGCGGCTGATTGTTGGACAACTAAAACGCCTTTGCCCAGGGCCAGAACTC-3’) oprF-3-1(5’-GAGTTCTGGCCCTGGGCAAAGGCGTTTTAGTTGTCCAACAATCAGCCGCAC-3’) and oprFint-3-2G (5’-TTATACGAGCCGGAAGCATAAATGTAAAGCAAGCTTGTGACCGTTGTCACGCTTCTC-3’).

The two PCR products were cloned into pEXG2 which had been digested with *Kpn*I and *Hin*dIII using NEBuilder® HiFi DNA Assembly (New England Biolabs). The resultant plasmid was verified by whole plasmid sequencing (Plasmidsaurus) and transformed into *E. coli* strain SM10l*pir.* The deletion construct was transferred into PA14 by mating, selecting for gentamicin resistance (30µg/mL gentamicin, 5µg/mL triclosan to select against the *E. coli* donor), and the resolved cointegrates were selected by sucrose selection at 30°C (5% sucrose LB agar plates with no salt). Sucrose resistant colonies were tested for loss of gentamicin resistance and screened for the presence of the deletion.

#### Complementation

Plasmid pMMB-*oprF* was generated by amplifying the *oprF* open reading frame from PA14 using primers 5GMMB (5’-TGAGCGGATAACAATTTCACACAGGAAACAGAATTCAGATGGGGATTTAACGGATG AAAC-3’) and oprF-3GMMB (5’-TGAAAATCTTCTCTCATCCGCCAAAACAGCCAAGCTTTTACTTGGCTTCAGCTTCTAC-3’). These were cloned into pMMB66EH (39) and cut with EcoRI and HindIII using the HiFi DNA Assembly Master Mix (NEB). Plasmid inserts were verified by DNA sequencing. PA14 and the Δ*oprF* mutants harboring either pMMB66EH(-) or pMMB-*oprF* were grown overnight in LB broth with 50µM IPTG and 150µg/mL carbenicillin. They were then diluted into high salt LB containing 5mM EGTA and 50µM IPTG and grown to late log phase.

### Western blot

Samples from each culture were normalized for optical density in a final volume of 75µL to which 25µL of 4x SDS sample buffer were added. Samples were then heated to 95°C for 10 minutes and separated on a 10% SDS PAGE gel, before being transferred to a PVDF membrane. The membrane was probed with an affinity-purified rabbit anti-ExoT antibody (40) at a dilution of 1:5000. and mouse anti-RpoA 4RA2 (BioLegend) at 1:10000. Anti-rabbit IrDye800 and anti-mouse IrDye680 antibodies (LiCor) were used as secondary antibodies (1:20000). Blots were scanned on a LiCor Odyssey M Imaging System.

### Corneal infection model

Males and females C57/Bl6/J mice (Jackson Labs) were anesthetized using ketamine/xylazine for corneal abrasion with 3 parallel 10 mm scratches with a 26½ gauge needle. *P. aeruginosa* strains were grown as described above and washed and resuspended in PBS to a concentration of 5 x 10^5^ CFU in 2 µL and applied topically to the abraded corneas. Mice were euthanized 24-48 hours after infection, and corneas were imaged, followed by eyes or corneas dissection. Animals that showed opacity in the contralateral eye or body injury before or after the infection were excluded from the study. All protocols were approved by UCI Institutional Animal Care and Use Committee under IACUC protocol AUP-21-123.

### Spatial transcriptomics workflow

For tissue fixation, 15 µL 4% formalin were dropped onto the eye of euthanized mice for 15 minutes. Eyes were then carefully resected and placed in 1 mL cold 4% at 4 °C for 48-72 hours. Eyes were placed in PBS and shipped to Moderna, where they were embedded by UMass Histopathology and processed by Umass Chan Medical School Morphology Core services using standard protocols.

Spatial transcriptomic experiments were performed as previously described using the BOND RXm (Leica) for antigen retrieval and the GeoMx DSP (Nanostring) for transcript collection and imaging (9). Probes hybridization was performed using Nanostring mouse whole transcriptome atlas (WTA) and custom *P. aeruginosa* probe sets overnight at 37 °C.

After collection, transcripts from each ROI were uniquely indexed by PCR amplification using i5 and i7 dual-indexing primers (Illumina). AMPure XP beads (Beckman Coulter) were used to purify the PCR product and concentration (Qubit fluorimeter, Thermo Fisher Scientific) and quality (TapeStation, Agilent Technologies) were assessed before sequencing.

### Data Processing and Quantification

The required sequencing depth was calculated by summing the illumination areas (in µm²) and multiplying the total by a factor of 100, following recommendations from the GeoMx DSP. Library and Sequencing Guide (NanoString Technologies). Sequencing was performed on the NovaSeq 6000 platform, generating FASTQ files that were converted to Digital Count Conversion (DCC) files using GeoMxNGSPipeline v.2.3.3.10, an open-source pipeline from NanoString Technologies. The DCC files were analyzed using NanoString’s GeoMx™ Digital Spatial Profiler. Key quality metrics, including raw read counts, alignment rate, and sequencing saturation per area of illumination, were carefully evaluated to assess library quality. Only data from regions of interest (ROIs) with more than 1 million raw reads, an alignment rate above 80%, and sequencing saturation exceeding 80% were included in downstream analysis. A “no template control” sample, lacking target probes, was incorporated during library preparation and sequencing to monitor for potential contamination.

Additionally, negative control probes with External RNA Controls Consortium (ERCC) sequences were included in each GeoMx™ RNA assay to estimate background noise and set a quantification threshold. Target probe data were filtered by frequency per ROI and normalized using background subtraction based on the highest negative control counts. Differential gene expression across experimental groups was identified using DESeq2 (41), a statistical tool for normalized gene expression analysis. Functional enrichment analysis was conducted with the Python library GOATools to interpret biological significance (42).

### Quantification of reactive oxygen species (ROS)

Bone marrow neutrophils were isolated from C57BL/6 (WT) mice and were enriched using EasySep™ Mouse Neutrophil Enrichment Kits (STEMCELL). Following enrichment, cells were concentrated at 1×10^6^ /ml in RPMI (Gibco) supplemented with 500µM luminol (Sigma Aldrich). Neutrophils were then added to black-sided optically clear 96-well tissue culture plates (Corning) at a concentration of 2×10^5^ per well. The ROS inhibitor diphenyleneiodonium chloride (DPI; Sigma Aldrich) was added to respective wells at a final concentration of 10µM and cells were incubated at 37°C, 5% CO2 for 30 minutes prior to stimuli being added. Bacteria were prepared (as described above) and concentrated to 1×10^8^ bacteria/ml (MOI 10 in 20µl) or 3×10^8^ bacteria/ml (MOI 30 in 20µl) and 20µl of PA14 parental, Δ*oprF or;*Δ*oprF:oprF* (complemented) strains were used to stimulate the cells. The protein kinase C activator phorbol 12-myristate 13-acetate (PMA; Sigma Aldrich) was at a concentration of 25nM was used as a positive control. Once stimuli were added, luminescence was measured every two minutes for 90 minutes using a BioTek Cytation5 (Agilent Technologies). Respective curves were plotted and the area under the curve was calculated using GraphPad Prism software.

### Quantification of corneal opacity

Corneal images were obtained using a Leica MZ10 F Modular Stereo Microscope with a Leica DFC450 C camera attachment. Image brightness was adjusted using ImageJ and kept consistent for all images. ROI selection excluded sclera, and mean intensities were background corrected by subtracting the mean intensity of naïve corneal images. Percent corneal intensity was determined as a ratio of corrected mean intensity to the intensity of a completely white cornea.

### Quantification of viable bacteria

Infected eyes were homogenized in 1 mL of sterile PBS using a 5 mm steel ball bearing at 30 Hz for 3 minutes using a Qiagen Tissue Lyser II. 10 uL of serially diluted homogenate were streaked onto LB agar plates and incubated at 37 C, 5% CO_2_ overnight. Colony forming units (CFU) were counted manually.

### Immunohistochemistry of corneal sections

For sections stained at UC Irvine, infected eyes were removed and fixed in Davidson’s Fixative (Polysciences) for 24hrs before transitioning to 70% ethanol. Eyes were then oriented in paraffin wax and sectioned into 8µm sections by the UC Irvine Ocular Microanatomy Core. Slides were deparaffinized and rehydrated to water with xylenes and an ethanol series respectively. For antigen retrieval, proteinase K (DAKO) was incubated 20 min at 37°C. Sections were then blocked with blocking buffer containing Fc block (BioLegend) and normal goat serum (Jackson ImmunoResearch) 1hr before staining overnight at 4°C with primary antibodies to Ly6G (NIMP-R14) and OprF (Thermo Fisher Scientific).

Sections were then washed and stained 1hr with secondary antibodies goat anti-Rat 647 and goat anti-rabbit 555 (Thermo Fisher Scientific). Sections were washed again before 5 minutes incubation with DAPI nuclear stain (Sigma Aldrich). Slides were mounted using Vectashield Antifade Mounting Media (Vector Laboratories), and sections were imaged on a Leica DMI6000 B Inverted Fluorescent Motorized Microscope, with Leica LAS-X software.

### Flow cytometry

Mice were infected using the corneal infection model described above. After 48h, mice were euthanized, and corneas were dissected and placed in dissociation buffer containing 3mg/ml collagenase (C0130; Sigma Aldrich) in RPMI (Gibco), 2% HEPES (Gibco), 2% sodium pyruvate (Gibco), 1% nonessential amino acids (Gibco), 1% penicillin-streptomycin (Gibco), 2.5% DNAse I (Qiagen), 10% heat-inactivated fetal bovine serum (FBS; Corning), and 0.5% calcium chloride (Sigma Aldrich). Samples were then incubated at 37°C water bath for 1hr. All steps following dissociation were done at 4°C. After dissociation, collagenase was neutralized using FBS and samples were filtered using 35µM filter Falcon™ tubes (Fisher Scientific).

Cells were centrifuged and resuspended in buffer containing 2% FBS and 1mM EDTA. Mouse Fc receptors were blocked for 5 min with anti-mouse CD16/32 Ab (BioLegend) and sections were stained for 40 minutes with fixable viability dye e780 (Invitrogen), CD45, Ly6G, CD11b and Ly6C (BioLegend). Sections were washed with staining buffer and fixed for 15 minutes using BD Biosciences Cytofix/Cytoperm™ and quantified using an ACEA Novocyte 3000 (Agilent Technologies) flow cytometer and NovoExpress software. Gating strategy is shown in Supplementary Figure 4.

### Statistics

For *in vivo* experiments, statistical significance was determined using ANOVA (GraphPad Prism). *In vitro* assays required a minimum of 3 biological repeats using the mean of four technical replicates. *P*< 0.05 was considered significant. The number of biological replicates for each experiment is noted in the figure legends.

## Author contributions

SA and MEM performed in vivo and in vitro experiments. HZ, NC, ON, AT and MG performed spatial transcriptomics and bioinformatics analyses. AR generated deletion mutants and complemented strains and performed western blots. SA, HZ, AR, EP and MG designed the experiments and wrote the manuscript.

